# Quantitative analysis of calcium oxalate monohydrate and dihydrate for elucidating the formation mechanism of calcium oxalate kidney stones

**DOI:** 10.1101/2022.10.20.513046

**Authors:** Mihoko Maruyama, Koichi P. Sawada, Yutaro Tanaka, Atsushi Okada, Koichi Momma, Masanori Nakamura, Ryota Mori, Yoshihiro Furukawa, Yuki Sugiura, Rie Tajiri, Kazumi Taguchi, Shuzo Hamamoto, Ryosuke Ando, Katsuo Tsukamoto, Kazufumi Takano, Masayuki Imanishi, Masashi Yoshimura, Takahiro Yasui, Yusuke Mori

## Abstract

We aimed to identify and quantitatively analyze calcium oxalate (CaOx) kidney stones on the order of micrometers, with a focus on the quantitative identification of calcium oxalate monohydrate (COM) and dihydrate (COD). Fourier Transform Infrared (FTIR) spectroscopy, powder X-ray diffraction (PXRD), and microfocus X-ray CT measurements (micro-CT) were performed, and the results were compared. The extended analysis method of the FTIR spectrum, focusing on the 780 cm^−1^ peak, made it possible to achieve a reliable analysis of the COM/COD ratio. We succeeded in the quantitative analysis of COM/COD in the region of 50 × 50 μm by microscopic FTIR for thin sections of kidney stones, and by the micro-CT for bulk samples. The extended analysis method of the FTIR spectrum focusing on the 780 cm^−1^ peak was introduced to analyze the COM/COD ratio. The analysis results based on PXRD measurement with micro sampling, microscopic FTIR analysis of a thin section, and micro-CT observation of a bulk sample of a kidney stone showed roughly consistent results, indicating that all methods can be used complementarily. This quantitative analysis method evaluates the detailed CaOx composition on the preserved stone surface and provides information on the stone formation processes and interactions with organic molecules.

## Introduction

Kidney stones contain ~90% mineral compounds and 10% organic compounds. The mineral compounds consist of calcium stones such as calcium oxalate (CaOx) and calcium phosphate, and non-calcium stones (e.g., struvite, uric acid, cystine, proteins, and drug stones) [1]. Kidney stone is a common disease, and approximately 1.7 – 14.8% of the population are affected once in their lifetime [2] [3]. Furthermore, the recurrence rate at 5 years was more severe than 40 %. Although the management of patients with symptomatic kidney stones has evolved from open surgical lithotomy to minimally invasive approaches such as percutaneous nephrolithotomy (PCNL), ureteroscopic lithotripsy (URSL), and shock wave lithotripsy (SWL), these are still invasive[4]. Considering the high occurrence and recurrence rate, understanding the pathogenesis of kidney stone formation and suggesting new endourological approaches that reduce the severe rate of kidney stone disease are essential [1, 5].

In many previous studies, the identification of crystal phases and proteins in kidney stones has been conducted to clarify the pathogenesis of the disease. In most cases, kidney stones are crushed and powdered. The crystal phase and organic substances contained in the kidney stones are identified using Fourier transform infrared (FTIR) spectroscopy, Raman spectroscopy, energy dispersive X-ray spectrometry (SEM-EDX), mass spectroscopy, etc. [6, 7]. However, identification using powdering samples had limitations; the spatial information of stones, in other words, the historical data of the stones, was lost. Identifying crystal phases and crystal texture classifications using slice sections and polished thin sections of kidney stones is an effective way to overcome this problem.

Observation of such thin sections of kidney stones by optical microscopy has been conducted using slice sections and polished thin sections of kidney stones for more than 70 years [8]. However, in those days, there were problems and limitations in the technology for shaping stones, composition analysis technology for minute regions, and complementary analysis technology for organic and inorganic substances; thus, the formation mechanism was not sufficiently clarified. For example, deterioration of the surface structure and protein expression occurs during the process of shaping kidney stone thin sections. Recently, the observation and analysis of kidney stone thin sections were focused on again, and we developed a new method for making thin sections without severe deterioration of the surface and protein expression. Using such thin sections of kidney stone, we succeeded to reveal micro-scale distributions of three different proteins using the advanced multiple immunofluorescence staining (multi-IF staining) [9]. The spatial distributions of these proteins in kidney stones are essential for evaluating the *in vivo* effects of proteins on stone formation. We need detailed mineral distribution in kidney stones to discuss the stone formation process and pathogenesis. Recently, high-resolution FTIR, Raman spectroscopy, etc., were introduced to observe thin sections. Castiglione *et al*. attempted chemical imaging of kidney stones using a confocal Raman microspectrophotometer and succeeded in identifying kidney stone components, including crystal polymorphisms of CaOx and calcium phosphates [10]. This method is helpful in determining the organization of components within stones but still has several limitations: identifying a kidney stone containing a high concentration of proteins is difficult because of its vast autofluorescence background, and quantification of each component is still an issue [10].

Quantification of CaOx phases by FTIR was discussed, and a reliable method was suggested for the bulk analysis of kidney stones [11, 12]. The KBr pellet technique and attenuated total reflection method (ATR method) enabled us to measure small amounts of powder samples; however, invasive sample preparation, for example, powdering a part of a kidney stone, was required. Blanco *et al*. reported the mapping of calcium oxalate monohydrate (COM), calcium oxalate dihydrate (COD), and carbonate apatite (CAP) by FTIR spectroscopy using kidney stone thin sections. The specific peaks of COM and COD (1630 cm^−1^ and 1680 cm^−1^ peaks) were focused on in this study for the quantification of COM and COD, but these peaks were affected by other phases such as phosphoric acid and organic molecules. Therefore, mapping the crystal phases based on these two peaks may lead to incorrect results. Yoshida et al. introduced a method for more reliable phase identification of COM and COD. This reference focused on the 780 cm^−1^ peak, which is relatively unaffected by other crystal components, and calculated their infrared absorption [13].

In this study, we aimed to identify and quantitatively analyze CaOx kidney stones in the order of micrometers. First, we targeted the most common calcium oxalate-based stones and focused on regional quantitative analysis of COM and COD. Kidney stones were collected from patients at Nagoya City University, and several thin sections and polished bulk samples were prepared. Polarized microscopy and FTIR were used for the crystal structure analysis and phase identification of the local points. The FTIR spectra were analyzed based on Ref. [13]. Powder X-ray diffraction (PXRD) measurements of a small amount of powder samples corrected from the thin section were conducted. The PXRD and FTIR results were compared to establish a reliable measurement procedure. We also analyzed a sliced bulk sample of a kidney stone using a microfocus X-ray CT system to quantify COM and COD. The quantification results of kidney stone samples using each method were consistent. The developed quantitative measurement procedures for CaOx polymorphs enabled us to investigate kidney stones in greater detail. The polymorphism of CaOx in vivo is crucial for understanding the stone formation process. Further investigation using quantitative measurement procedures of CaOx combined with the visualization of multiple proteins in human kidney stones[9] will elucidate the kidney stone formation processes.

## Materials and Method

### Ethics Statement

The research project presented in this paper was approved by the institutional review board of the Graduate School of Medicine, Nagoya City University. All methods were carried out in accordance with the relevant guidelines and regulations. Written informed consent was obtained from all subjects according to the procedures approved by the ethical committee board.

### Stone collection and preparation of stone sections

CaOx kidney stones were selected from thousands of human kidney stone collections from the Department of Nephro-Urology, Nagoya City University, Japan. Sample No. of the stone was 1004. Sample 1004 was partly crushed and powdered for bulk FTIR analysis. Thin sections were prepared using the resin-embedding method. The kidney stone was wholly embedded in epoxy resin CaldoFix (Struers). We manually polished the sample on glass using an abrasive (Al2O3). After each grinding and polishing step, the sections were washed to remove abrasives. A sample with 2 mm thickness was prepared for the microfocus X-ray CT system (inspeXio SMX-100CT). The remaining piece was affixed to a glass slide using epoxy adhesive Bond-E (Konishi), with the polished surface facing down. Subsequently, we made a second cut parallel to the glass slide with a thickness of approximately 1 mm, and the sample was ground and polished to a thickness of 20-30 μm. It was necessary to obtain a smoother surface for chemical analysis, such as the FTIR reflection method; therefore, we polished it again using a diamond slurry. In this study, we prepared two thin sections of the kidney stone [14].

### Observation by a polarized-light microscope

We observed the size, coloration, and extinction of crystals in the kidney stone thin section using a polarized-light microscope (Nikon, OPTIPHOT2-POL), switching between open Nicol and cross Nicol.

### Measurement by microscopic FTIR

We analyzed Sample1004 thin section using microscopic FTIR (JASCO, FT/IR-6100). The reflection method was used for measurement. The settings were as follows: cumulative number, 256; measurement range, 600 ~ 4000 cm^−1^, measurement area, 50×50 μm. Because the obtained specular reflection spectrum was affected by the anomalous dispersion of the refractive index, we performed the Kramers-Kronig transformation after removing the absorption of H_2_O and CO_2_ in the atmosphere.

### Quantitative analysis of COM and COD using the FTIR spectrums

Ref. [13] shows the quantitative analysis method of powdered COM and COD using FTIR. In our study, we attempted to extend this method to analyze kidney stones. We measured the prepared powder samples (shown in the supplemental material) using microscopic FTIR spectroscopy (JASCO, FT/IR-6100). A transmission method was used for the measurement. The settings were as follows: cumulative number, 256; measurement range, 600 ~ 4000 cm^−1^, measurement area, 50×50 μm. The absorptions of H_2_O and CO_2_ were removed from the obtained spectra to eliminate atmospheric influence. The measurement range included the O-H stretching bond at 3200–3550 cm^−1^, C=O stretching bond around 1700 cm^−1^, C-O stretching bond around 1300 cm^−1^, and C-H bending bond at 675–900 cm^−1^ (Fig. 1 (a)) [15].

**Figure 1.**
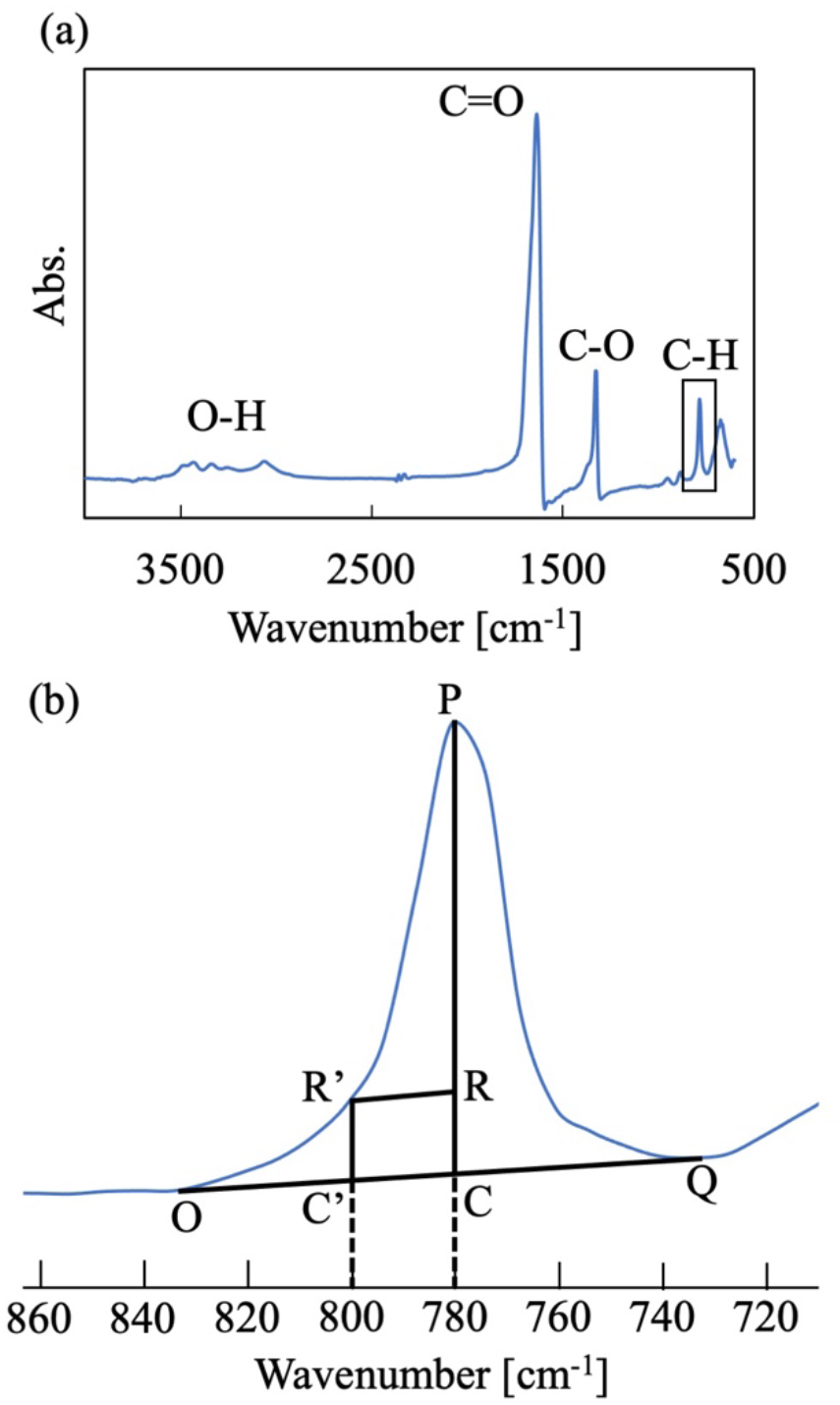
(a) The IR spectrum of calcium oxalate kidney stones. (b) The IR spectrum focused on a peak at 780 cm^−1^. Point P is the peak point at 780 cm^−1^. Point Rʼ is the measurement value at 800 cm^−1^. Segment OQ is the line connecting the bottoms of both valleys of the peak. Segment R’R is the parallel line with segment OQ. Segment PR shows the amount of absorption contributed by COM, and segment RC offers the amount of absorption contributed by COD.

We mainly focused on the range of 700 to 900 cm^−1^. Although the infrared absorption of COM has a peak at 780 cm^−1^ and that of COD also has a peak at 780 cm^−1^, the infrared absorption of COD has a larger half-maximum width than that of COM. Thus, COD’s 780 cm-1 absorption peak could be fully differentiated from that of COM at approximately 800 cm^−1^. Here, we provide a summary of the analysis method in Ref. [13]. First, we calculated the base value of the 780 cm^−1^ peak (point C) by drawing a line (segment OQ) connecting the bottoms of both valleys of the peak (Fig. 1 (b)). Next, we drew a parallel line (segment R’R) with segment OQ from the measured value at 800 cm^−1^ (point Rʼ). The vertices of lines OQ and PC are defined as points R. The ratio of line PR to line PC corresponds to the absorption contribution of COM at the 780 cm^−1^ peak, and the ratio of line RC to line PC corresponds to that of COD at the 780 cm^−1^ peak. We defined the absorption ratio of COM to total calcium oxalate as *M*, and that of COD as *D*. We also described the proportion of COM as *m* and COD as *d* for the calcium oxalate samples. Here, they can be approximated by the following linear equations, where *a*, *b*, *a’,* and *b*ʼ depend on the apparatus used: 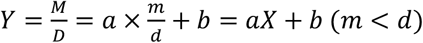

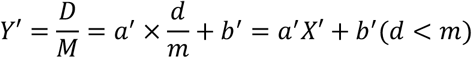

To determine *a*, *b*, *a’,* and *b*ʼ for the apparatus used (JASCO, FT/IR-6100), we drew a calibration curve using prepared standard samples. COM and COD were mixed at arbitrary molar ratios (see the supplemental). Each of the collected standard samples was measured more than six times. Each measured IR spectrum was treated as previously described. We measured the infrared absorption of the kidney stone samples and calculated the content ratio of COM and COD using this calibration curve.

### Measurement by powder X-ray diffraction

We identified the crystal components in the kidney stones by local analysis using PXRD. After optical observation, we scraped off part of the thin section and measured the samples by PXRD. The scraped areas are shown in Fig. 2 (a) by black arrows (approximately 50×50 μm). PXRD data were measured at room temperature on a Rigaku SmartLab powder diffractometer (45 kV, 200mA, rotating anode) in the Debye–Scherrer geometry with Cu Kα_1_ radiation monochromatized by a Johansson Ge crystal and focused by a multilayer mirror. Region B in Fig.2 (b) was measured using Mo Kα_1_ radiation. The PXRD patterns for COM and COD were calculated using VESTA with references [16] and [17].

**Figure 2.**
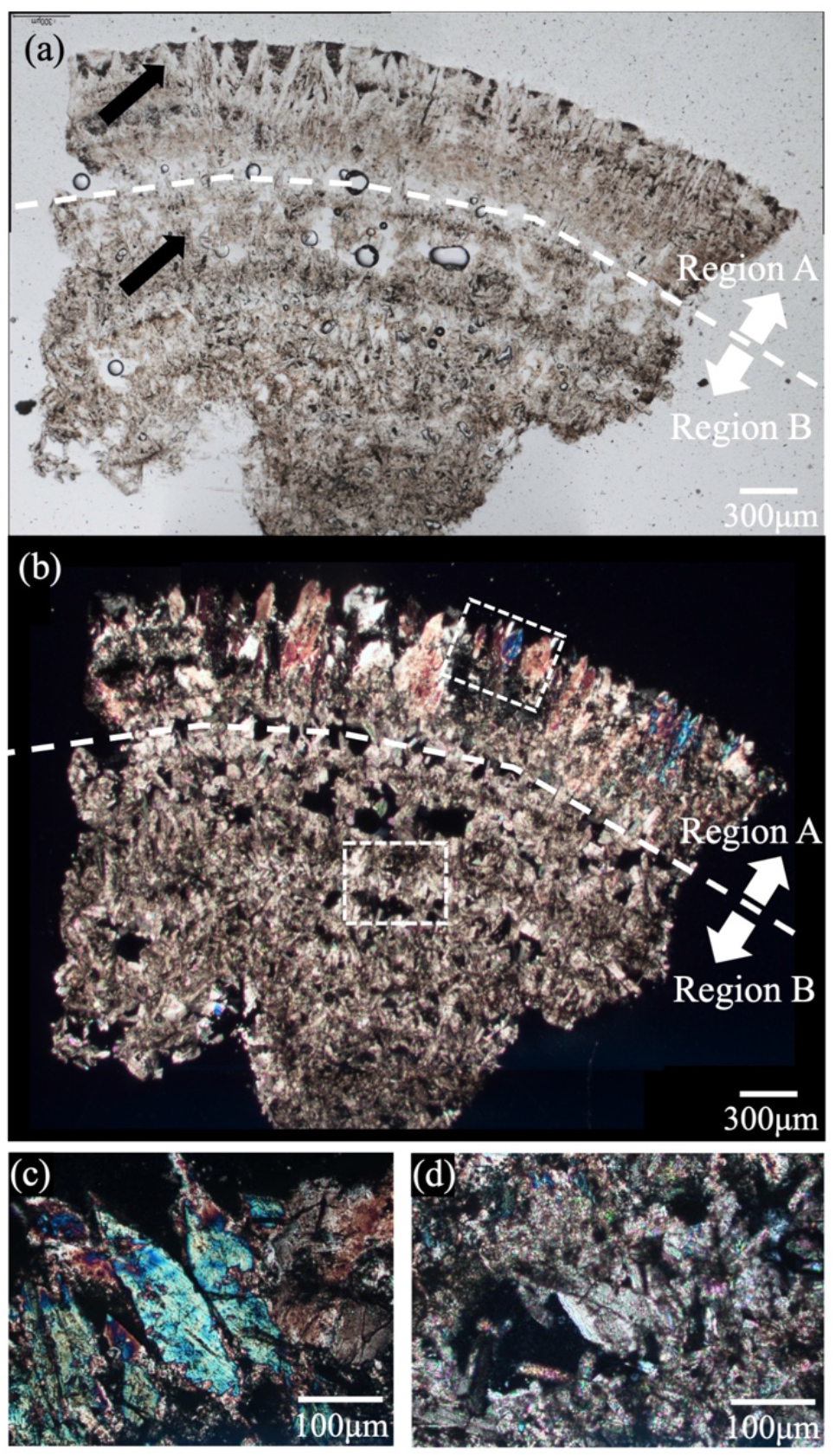
(a) and (b) show open-Nicol images and cross-Nicol images of sample1004 observed using a polarization microscope. (c) and (d) show enlarged images of regions A and B observed using a polarization microscope (inside the dashed square in (b)).

### Measurement by microfocus X-ray CT system

We identified the crystal components in the kidney stones using a microfocus X-ray CT system (inspeXio SMX-100CT Plus). The measurement parameters were X-ray tube (90 kV, 44 μA) and voxel size (0.005 mm/voxel). We measured a 2 mm-thick kidney stone sample and obtained a series of X-ray tomographic images. The main components of CaOx kidney stones, COM and COD, exhibit different amounts of X-ray absorption [18]. Approximate local phase identification was performed based on the differences in the values. The analysis procedure using the obtained X-ray tomographic images is shown in the supplemental information.

## Results

Figures 2 (a) and (b) show images of sample1004 thin section observed by a polarization microscope; (a) is an open-Nicol image and (b) is a cross-Nicol image. The color of the crystals that make up the kidney stones ranges from light brown to brown. The color distribution was seen concentrically in the stone. The thinness of the sample (approximately 20–30 μm) and surface smoothness made it possible to observe kidney stone samples clearly by optical methods and to analyze them by the FTIR reflection method. We divided the thin section into two areas (regions A and B), as shown in Fig. 2 (a) and (b), because the structures and crystal sizes were different. These areas were observed and analyzed separately using various methods. The details of this process are presented below.

Figure 2 (c) and (d) show magnified cross-Nicol images of the crystals in regions A and B, respectively. Region A was mainly composed of crystals with sizes of the order of 100 μm, and the colors varied from blue to magenta with a thickness of 20-30 μm. The color tone faded as the observation stage was rotated and then disappeared entirely at an extinction angle of approximately 45°. With further rotation, the crystals’ brightness faded from about 45° and showed maximum intensity when the rotation angle was approximately 90°. The optical behavior indicated that large crystals (100 µm in size) were almost single crystals. Region B was composed of crystals with dimensions of the order of 10 μm, and the color was iridescent with a thickness of 20-30 μm because of internal interference. Determining the extinction angle of each crystal is difficult because of the iridescent color.

We performed local sampling from the thin section of the sample for the phase identification of small regions using PXRD. Figure 3 shows the PXRD patterns of Regions A and B. The PXRD pattern of region A (Fig.3 (a)) showed that there were specific peaks of COD (peaks at approximately 14.3°, 20.1°, and 32.2°) and no specific peaks of COM (peaks at about 14.9°, 24.4°, and 30.1°). The PXRD pattern of region B (Fig.3 (b)) shows specific peaks of COM, and no specific peaks of COD were detected. These results concluded that region A was mainly composed of COD crystals, and region B was mainly composed of COM crystals.

**Figure 3.**
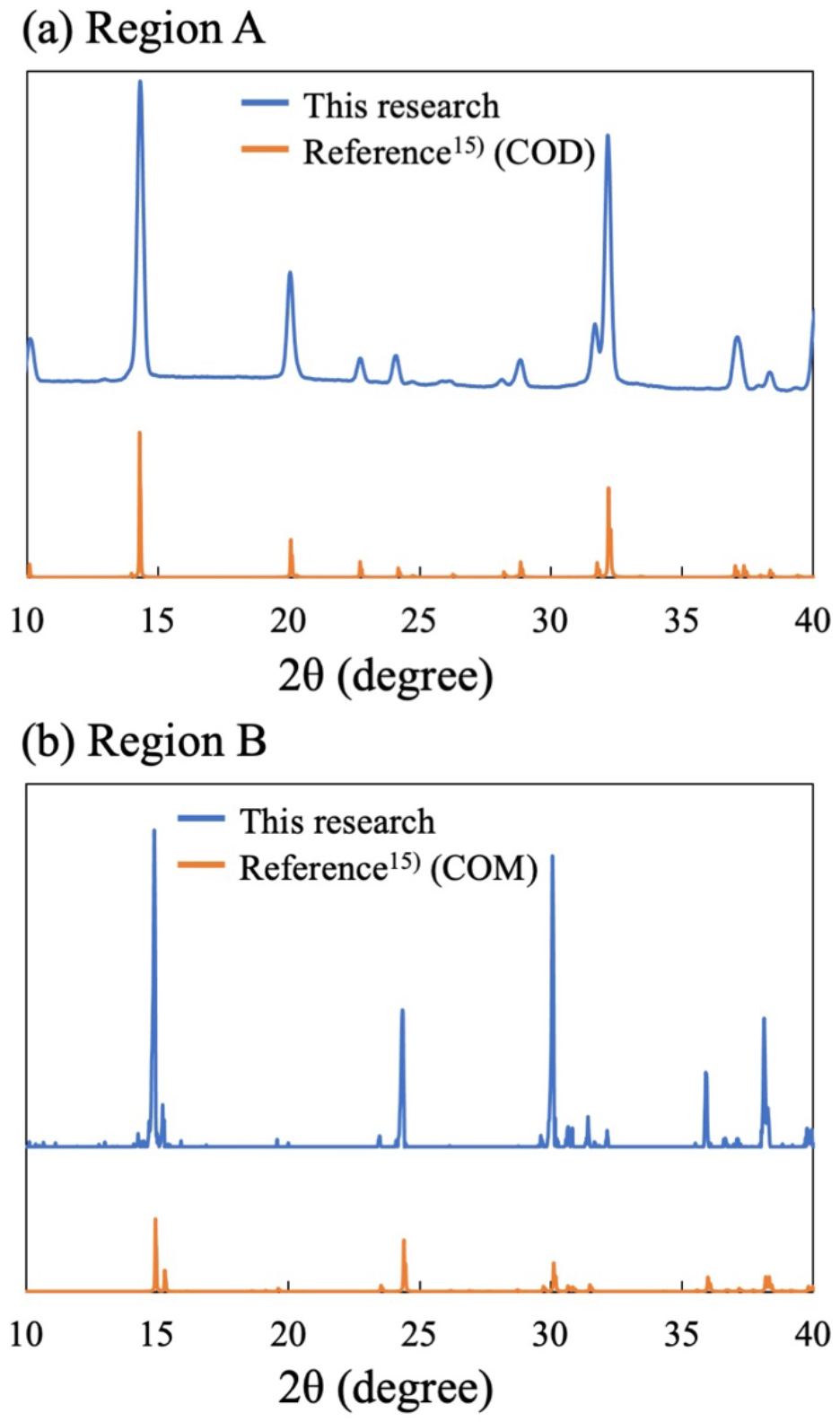
The measurement result of Sample1004 by PXRD. (a) shows the result of Region A and (b) shows that of Region B in Fig. 2. Each upper graph is the result of this research, and each lower graph is that of COM or COD by Ref. [19]

Figure 4 (a) shows an open-Nicol image of the thin section of sample1004. Note that the thin section was the second shaped from the same kidney stone (Sample1004). Although the shape is different from that shown in Fig. 2, the second thin section was also divided into two regions (A and B) based on observations using a polarized-light microscope. We indicated the FI-IR measurement points in the image by solid squares with numbers (from *1* to *4*). Figure 4 (b) shows a set of IR spectra at each measurement point. The peaks of calcium oxalate, as shown in Fig. 1, were also identified in each spectrum. The absorption features derived from the O-H band (3200–3550 cm^−1^) and absorption on the lower wavenumbers (C=O stretching bond around 1700 cm^−1^, C-O stretching bond around 1300 cm^−1^, and C-H bending bond at 675–900 cm^−1^) indicated that the main phase at measurement points *1* and *2* were COD and that at measurement points *3* and *4* were COM [15]. To obtain more details on the ratio of COM to COD, we analyzed the spectrum focusing on the peak near 780 cm^−1^ (Analysis 1). We confirmed that the COD ratio was 100±10 wt% at measurement points *1* and *2*. This proves that the main CaOx phase in region A is COD. In contrast, the COM ratio at measurement point *4* was 100±10 wt%. This shows that the main CaOx phase in region B is COM. At the boundary between regions A and B, similar to measurement point *3*, the crystal phases observed in regions A and B were mixed. According to our calculations, the ratio of COM was estimated to be 70±10 wt%, and that of COD was estimated to be 30±10 wt% from the IR spectrum. For comparison, we analyzed the FTIR spectra using the methods described in Ref [20] (Analysis 2) and Ref [11] (Analysis 3). Martin *et al*. (Ref [20]) focused on the absorption bands at 910 cm^−1^ and 780 cm^−1^. Mauric-Estepa (Ref [11]) focused on the peak shift of the absorption at 1324 cm^−1^. The results calculated using these three methods are summarized in Table 1. The results of Analysis 2 tended to indicate a lower COD amount compared to the results of Analysis 1. The results of analysis 3 showed that the ratio of COM to COD was approximately half at each 1–4 point. PXRD results showed that region A (containing measurement areas 1 and 2) is mainly composed of COD, contrary to region B (containing measurement point 4), composed of COM. The results of analysis 2 showed that measurement areas 1 and 2 contained 50 ~ 57% COM, and analysis 3 also showed 50 ~ 63% COM, but they were not consistent with the PXRD results. The results of Analysis 1 matched well with the PXRD dataset.

**Figure 4.**
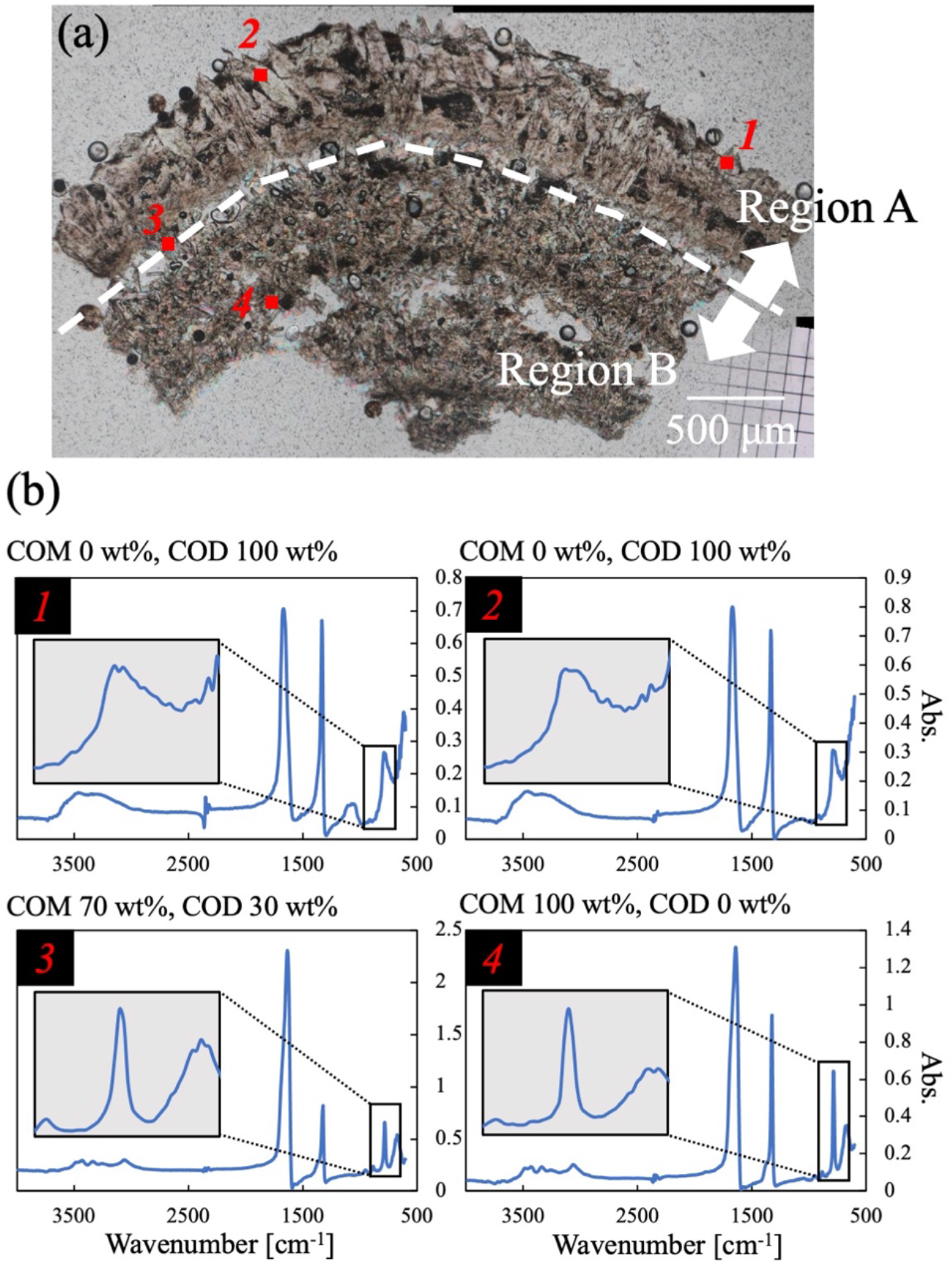
(a) shows an open-Nicol image of the second section of Sample 1004. Red marks and numbers indicate the FTIR measurement points. (b) shows the measurement results for each measurement point obtained by microscopic FTIR. The inner graphs are the enlarged views of the peaks at 780 cm^−1^ in each graph, and the ratio of COM to COD in the upper part of the graph shows the ratio calculated by the analytical method derived in this study.

**Table 1.** The ratio of COM and COD calculated by three different analyses.

|  | Analysis 1 Ref. [13])<br>(780 cm<sup>-1</sup>) |  | Analysis 2 (Ref. [19])<br>(910 and 780 cm<sup>-1</sup>) |  | Analysis 3 (Ref. [11])<br>(1324 cm<sup>-1</sup>) |  | PXRD |
| --- | --- | --- | --- | --- | --- | --- | --- |
|  | COM | COD | COM | COD | COM | COD |  |
| ① | 0% | 100% | 50% | 50% | 50% | 50% | COD |
| ② | 0% | 100% | 57% | 43% | 63% | 47% | COD |
| ③ | 70% | 30% | 100% | 0% | 48% | 52% | - |
| ④ | 100% | 0% | 100% | 0% | 54% | 46% | COM |

Figure 5 shows a series of tomography images of sample 1004 obtained using a microfocus X-ray CT system (inspeXio SMX-100CT Plus). To get more details on the ratio of COM to COD, we analyzed 11 measurement points, as shown in Fig.5. Each measurement point is shown as a solid red square, and each of them is 50×50 μm. Calculations showed that the ratio of COD of measurement points in Region A (10, 11) was more than 90%. The percentage of COD of measurement points in Region B (1, 2, 3, 4, 6, 7, 8) was less than 10% (in other words, the ratio of COM was more than 90%), except for points 7 and 8. The percentages of COM and COD at points 7 and 8 were (80%, 20%) for 7 and (88%, 12%) for 8. The ratios of COM and COD at points 5 and 9, which were located between region A and region, were (83%, 17%) for 5 and (85%, 15%) for 9. Points 7 and 8 were also located near the boundary areas of regions A and B; thus, they exhibited relatively high COD rates. The observation and analysis results of the microfocus X-ray CT system were almost consistent with the results obtained by PXRD and microscopic FTIR (analysis 1). In particular, we can get a sequence of similar CT images in the measurement of the microfocus X-ray CT system. Thus, by performing the same analysis on each image of a sequence, it is also possible to calculate the total ratio of COM and COD contained in a non-destructive kidney stone sample.

**Figure 5.**
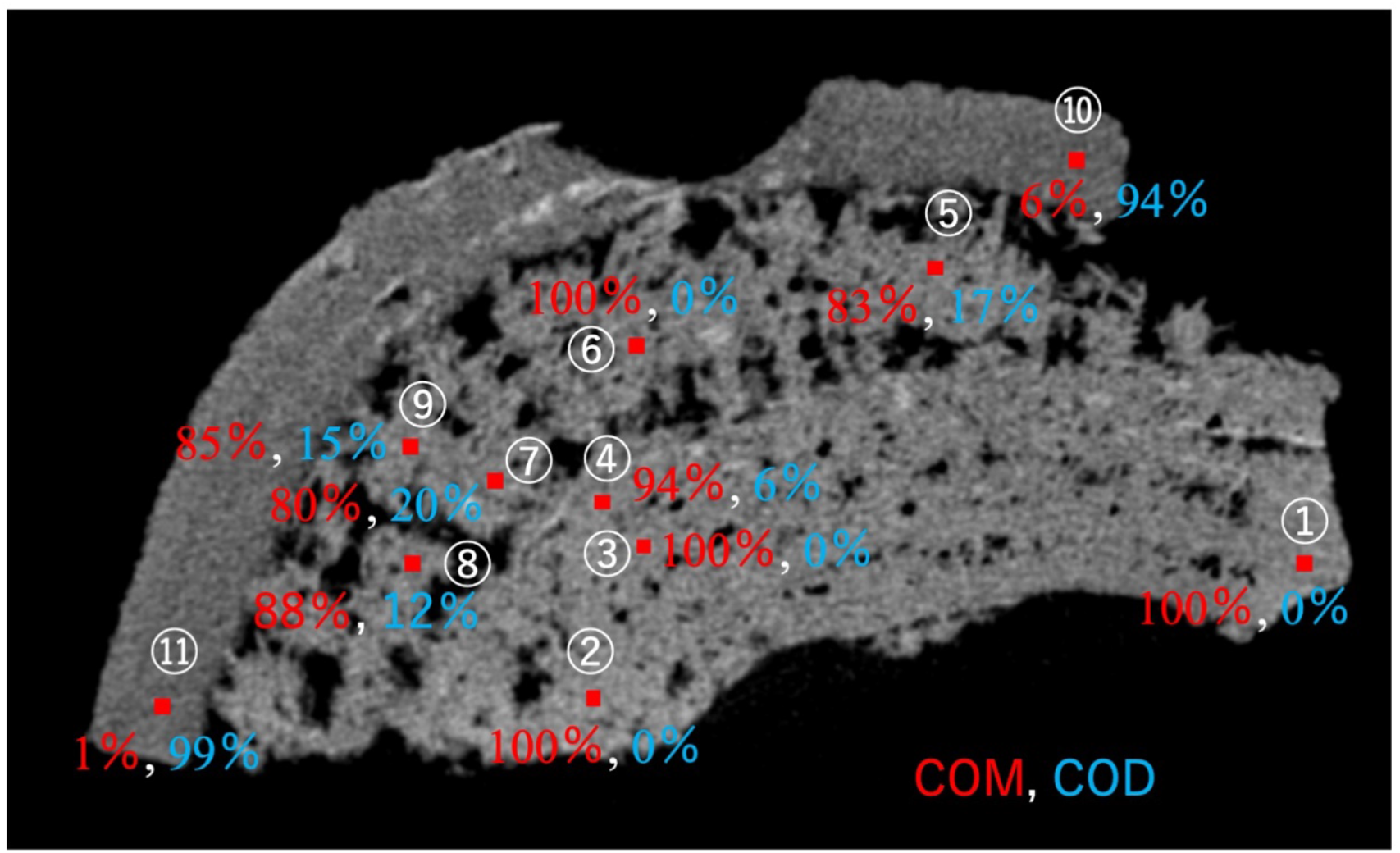
An image of microfocus X-Ray CT. Eleven analysis points are indicated with red solid squares and numbers. The analyzed results (the ratio of COM and COD in 50×50 mm square) are also shown. Red numbers correspond to the COM ratio, and blue numbers correspond to the COD ratio for each area.

As described above, we succeeded in the quantitative analysis of COM/COD in the region of 50 × 50 μm by microscopic FTIR for thin sections of kidney stones and by the microfocus X-ray CT system for bulk samples of kidney stones. The analysis results based on PXRD measurement with micro sampling, microscopic FTIR analysis of a thin section, and microfocus X-ray CT system observation of a bulk sample of a kidney stone showed roughly consistent results, indicating that all methods can be used complementarily. By adequately selecting these methods, considering the sample shape and/or sample processing, it will be possible to study kidney stones with higher accuracy than in previous studies.

## Discussion

We considered the formation mechanism of Sample1004 from the above results. From the cross-Nicol image in Fig. 2 (b), it was found that region B had many voids. The stone’s central part (region B) was mainly composed of COM, and the outer part (region A) was mainly composed of COD. Generally, the center part of a stone is the oldest and the outer part is the newest, as tree rings. At first, we assumed that region B of the stone was older, and region A was more recent. However, we have to consider the phase transformation. Sivaguru et al. reported that COD dissolves and transforms into COM in the human body[21]. It was also said that when the phase transformation from COD to COM occurs, a volume change occurs by discharging water occured[21]. If COM crystals nucleated and gathered each other in the early stage of stone formation, it is difficult for COD crystals to nucleate on the surface of COM crystals because of the solubility and supersaturation difference between COM and COD. In the internal environment of humans, the solubility of COD is higher than that of COM; it means that the supersaturation of COM is always higher than COD[22]. Thus more acceptable stone formation hypothesis is as follows; COD crystals nucleated and grew in the early stage of stone formation, then COD gradually transformed into COM from the center part of the COD stone. The previous nucleation of COD is a natural phenomenon because when supersaturation is sufficiently high for both the stable phase and metastable phase, the metastable phase can nucleate preferentially because of its lower interface energy (Ostwald’s step rule) [23]. In other materials, similar phenomena often occur[24, 25]. The transformation from COD to COM is mainly solution-mediated transformation[21]; dissolutions of COD crystals are needed. However, supersaturations of COM and COD in urine are always high enough for CaOx crystals to grow. Where can COD dissolve in urine? It is the inside of a kidney stone. The surface of a stone is constantly exposed to urine, which is a highly supersaturated environment; thus, there is no chance of dissolving COD and COM. In the case of a COD kidney stone, many COD crystals gather and the stone has many gaps between crystals. The center of the stone becomes a semi-closed system. Many gaps between crystals let urine enter inside of the stone where COD crystals can continue to grow using Ca^2+^, C_2_O_4_^2−^ and other solutes. The flow inside of the kidney stone is stagnated because of the narrow pass between crystals; thus sometimes the supply of solutes becomes not enough. If there are only COD crystals, crystal growth stops when the supersaturation of COD reaches nearly the equilibrium point. However, in most cases, COM, the stable phase of CaOx in the internal environment of the human, nucleate following COD crystal nucleation and growth. If the supersaturation of COD is near equilibrium and COM crystals co-exist, COM can still grow and Ca^2+^ and C_2_O_4_^2−^ consumed, then finally the solution will reach undersaturated for COD crystals. At the area, COD will partially dissolve, and Ca^2+^ and C_2_O_4_^2−^ will be released. The released solutes will be immediately consumed by COM near the dissolving COD. By this cycle, the transformation from COD to COM will gradually proceed. The solution-mediated transformation from a metastable phase to the stable phase often occurs in other materials[26–28]. Finally, the mosaic texture composed of irregular-oriented COM crystals appears from the center part of the kidney stone (Fig2. (d)). The transformation stops or significantly slows down where the supply of solutes from urine and the consumption of solutes by COM growth are balanced. In our sample, the boundary between regions A and B were probably such areas (Fig2. (a) and (b), and Fig.5)). The supply of solutes for COD and COM is enough outside of the boundary; thus we often observe kidney stones that have COM core area and COD rim area[15].

CaOx in kidney stones partially dissolves in the human body[21], but the dissolution process is followed by phase transformation from COD (a metastable phase of CaOx) to COM (the stable phase of CaOx). Through the transformation, a kidney stone becomes more durable, and also its structure probably becomes tight and rigid in the human body. This is a severe process in kidney stone pathogenesis; thus further research for understanding the mechanism of kidney stone formation is needed. The final structure of kidney stones and/or phase transformation speed estimated from the structure suggests the history of kidney stone growth and urine environment changes. Combined with our recent technique, multicolor imaging of calcium-binding proteins in human kidney stones [9], we can evaluate various kidney stone samples, and the information that stones have will reveal the factors of the stone formation processes and the interaction with organic molecules.

## Conclusion

In this study, we intended to perform phase identification and quantitative analysis of CaOx kidney stones in the micrometer order maintaining their spatial information. Polarized microscopy, FTIR, and PXRD measurements were performed for the crystal structure analysis and phase identification of local points using a thin section. We also conducted a quantitative analysis of CaOx in a sliced bulk sample of a kidney stone using a microfocus X-ray CT system. The extended analysis method of the FTIR spectrum, focusing on the 780 cm^−1^ peak, made it possible to perform a reliable analysis of the COM/COD ratio. Using polarized microscopy, we mapped the two-dimensional distribution of crystal phases with a reliable COM/COD ratio. Furthermore, the data from a microfocus X-ray CT system also gave us a reliable quantitative COM/COD ratio using the bulk sample of a kidney stone. This method enables us to discuss in more detail the location where crystal nucleation and phase transition occurs, how such processes advance, and the environmental changes during the processes. The polymorphism of CaOx in vivo is crucial for understanding the stone formation process. Further investigation using quantitative measurement procedures of CaOx combined with the visualization of multiple proteins in human kidney stones[9] will elucidate the kidney stone formation processes. We believe that this new methodology, examining the crystal phase and quantifying CaOx, could help understand the time course of stone formation with environmental changes that accelerate the process.

## Supporting information

shown in the supplemental material

## Acknowledgments

We thank K. Murakami, H. Kubo, and K. Kawamura for their support with the FTIR analysis. We also thank the members of the Medical and Engineering Tactics for Elimination of Rocks (METEOR) Project for their helpful discussions. Part of this study was supported by the JSPS KAKENHI Grant-in-Aid for Challenging Exploratory Research (No. 19K22965 and 20K21658) and research fellowships of JSPS (No. 18J40134). This work was also supported by the Konica Minolta Science and Technology Foundation, Shiseido Female Researcher Science Grant, Caterpillar STEM award 2019, and the Osaka University Program for the Support of Networking among Present and Future Researchers for M. M. This work was supported in part by Grants-in-Aid for Scientific Research from the Ministry of Education, Culture, Sports, Science, and Technology, Japan (No. 20K21658), the Naito Foundation, and the Hori Sciences and Arts Foundation.

## Author contribution statement

M. M. designed the study and wrote the manuscript. K. P. S., R. M., and M. M. carried out crystal phase identification using polarized microscopy, PXRD, and FTIR with the help of K. M.. K. P. S. and M. M. performed the microfocus X-ray CT with the help of M. N.. R. T. prepared all stone sections. All the authors discussed the results and critically revised the manuscript for important intellectual content.

## Competing Interest Statement

The authors declare no competing interests.

## Notes

### Competing Interest Statement

The authors have declared no competing interest.

