## Supplementary material for "Quantitative analysis of calcium oxalate monohydrate and dihydrate for elucidating the formation mechanism of calcium oxalate kidney stones": shown in the supplemental material

### Preparation of standard samples for creating an FTIR calibration curve

We prepared a standard powder sample of COD crystal for the calibration curve of FT-IR. The following reagents were purchased from FUJIFILM Wako Pure Chemical Corporation: calcium oxalate monohydrate (COM) (039-00752), sodium oxalate (198-02655), calcium chloride (038-12775). Filter papers were purchased from ASFIL: Membrane Filter (025045-MFPVDF). Ultrapure water used in the experiment was refined by Komatsu Electronics UL Pure (18.2 MΩ cm).

Calcium oxalate dihydrate (COD) powder used as a standard sample in this paper was synthesized by the method reported by Ref. [1]. First, 300 ml of sodium oxalate solution in ultrapure water (0.005 M) at room temperature was added to 500 ml of calcium chloride solution in ultrapure water (1 M) at 4℃ in a glass tube (1 cm diameter, 15 cm height). The mixture was left without agitation for 24 hours at 4℃. The deposited crystals were separated by filtration (ASFIL, Membrane Filter, 025045-MFPVDF) and characterized by powder X-ray diffraction. PXRD data of the COD powder were measured at room temperature on a Rigaku SmartLab powder diffractometer (45kV, 200mA, rotating anode) in Debye–Scherrer geometry with Cu $K\alpha_1$ radiation monochromatized by a Johansson Ge crystal and focused by a multilayer mirror. We carried out Rietveld analysis by using Rietan-FP[2] with initial structural models adopted from references [3, 4]. The refinement converged to $R_{wp}$ = 3.662 %, $R_{exp}$ = 2.697 %, S = $R_{wp}/R_{exp}$ = 1.3581. The refined mass fractions are 90.5 wt% of COD and 9.5 wt% of COM. Agreement factors for each phase are $R_B$ = 4.09 % and $R_F$ = 2.815 % for COD, and $R_B$ = 12.340 % and $R_F$ = 4.267 % for COM. The X-ray diffraction pattern is presented in Fig. S1.

Next, we prepared standard samples for FT-IR measurement calibration. COD powder obtained by the former method and commercially purchased COM powder were mixed at arbitrary ratios. At that time, the powders were carefully mixed without excessive force with a mortar to prevent the phase transformation during mixing. The ratios of COM in the mixture were set as 19 %, 32 %, 46 %, 64 %, 77 %.

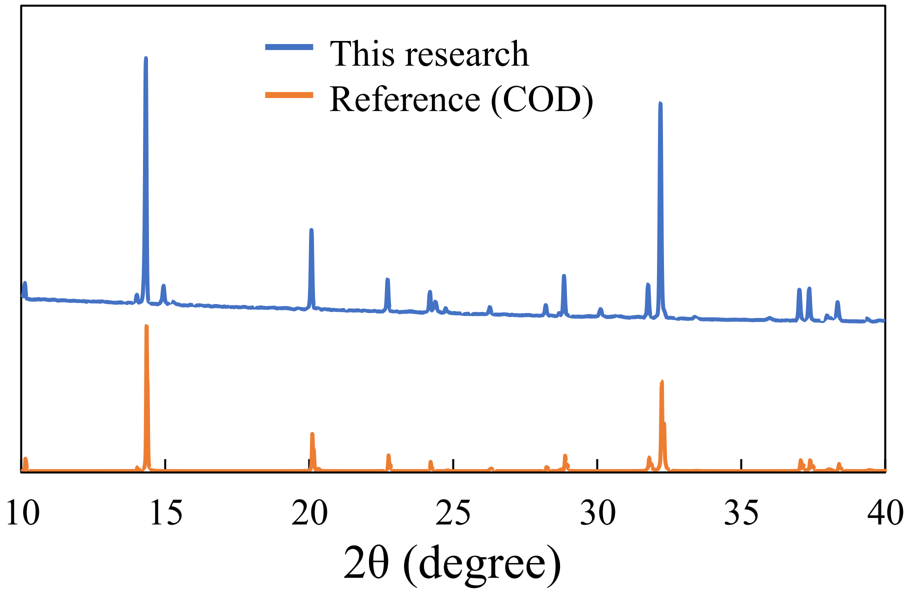


Figure S1 PXRD pattern of COD powder used as a standard sample by PXRD. The upper graph is the result of this research and the lower graph is that of COD by Ref. [3].

**Quantitative analysis of microfocus X-Ray CT images**

The brightness values in the region where COM and COD are 100% in the stone (indicated as red and blue circles in Fig. S2) were measured using image analysis software (ImageJ). These values were used as the reference values for COM and COD. Next, determined a measurement area of 50 μm square, then counted the number of pixels corresponding to the COM reference value, COD reference value, and others. For example, the measurement points shown in Figure 5 were confirmed to be the reference value of COM were 65 points, COD were 25 points and others were 10 points. Based on the calculated score, calculate the ratio of COM, COD, and other components in that area. The software developed independently was used to count the number of pixels.

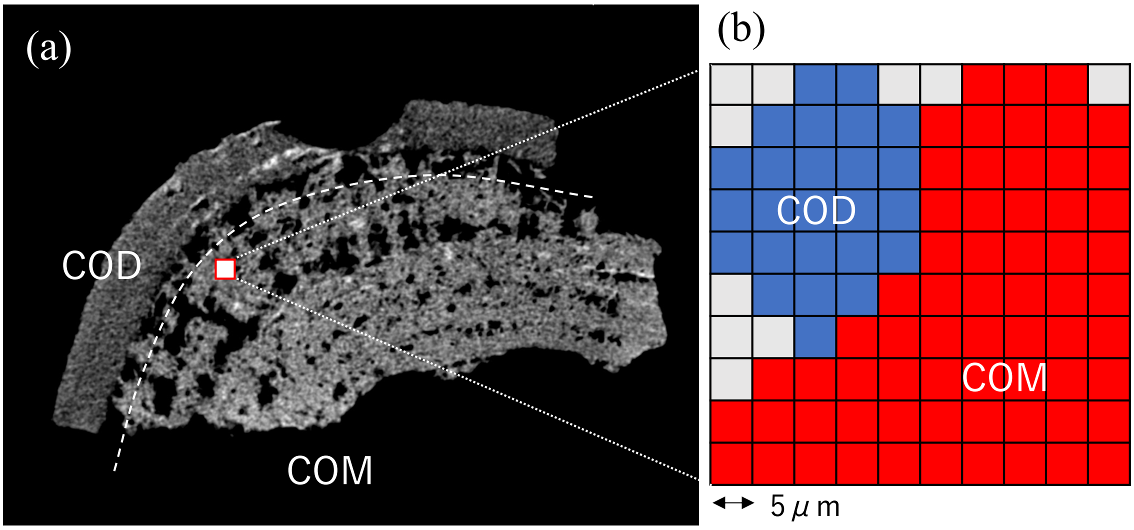


Figure S2 A microfocus X-ray CT image and the analysis procedure.
